# Comparative genomics shows that viral integrations are abundant and express piRNAs in the arboviral vectors *Aedes aegypti* and *Aedes albopictus*

**DOI:** 10.1101/128637

**Authors:** Umberto Palatini, Pascal Miesen, Rebeca Carballar-Lejarazu, Lino Ometto, Ettore Rizzo, Zhijian Tu, Ronald van Rij, Mariangela Bonizzoni

## Abstract

**Background:** Arthropod-borne viruses (arboviruses) transmitted by mosquito vectors cause many important emerging or resurging infectious diseases in humans including dengue, chikungunya and Zika. Understanding the co-evolutionary processes among viruses and vectors is essential for the development of novel transmission-blocking strategies. Arboviruses form episomal viral DNA fragments upon infection of mosquito cells and adults. Additionally, sequences from insect-specific viruses and arboviruses have been found integrated into mosquito genomes.

**Results:** We used a bioinformatic approach to analyze the presence, abundance, distribution, and transcriptional activity of integrations from 425 non-retroviral viruses, including 133 arboviruses, across the presently available 22 mosquito genome sequences. Large differences in abundance and types of viral integrations were observed in mosquito species from the same region. Viral integrations are unexpectedly abundant in the arboviral vector species *Aedes aegypti* and *Ae. albopictus*, but are ∼10-fold less abundant in all other mosquitoes analysed. Additionally, viral integrations are enriched in piRNA clusters of both the *Ae. aegypti* and *Ae. albopictus* genomes and, accordingly, they express piRNAs, but not siRNAs.

**Conclusions:** Differences in number of viral integrations in the genomes of mosquito species from the same geographic area support the conclusion that integrations of viral sequences is not dependent on viral exposure, but that lineage-specific interactions exits. Viral integrations are abundant in *Ae. aegypti* and *Ae. albopictus*, and represent a thus far unappreciated component of their genomes. Additionally, the genome locations of viral integrations and their production of piRNAs indicate a functional link between viral integrations and the piRNA pathway. These results greatly expand the breadth and complexity of small RNA-mediated regulation and suggest a role for viral integrations in antiviral defense in these two mosquito species.

## BACKGROUND

Nearly one-quarter of emerging or resurging infectious diseases in humans are vector-borne [1]. Hematophagous mosquitoes of the *Culicidae* family are the most serious vectors in terms of their worldwide geographic distribution and the public health impact of the pathogens they transmit. The *Culicidae* is a large family whose members separated between 180 to 257 Mya into the *Culicinae* and *Anophelinae* subfamilies [2]. Mosquitoes of the *Aedes* and *Culex* genera within the *Culicinae* subfamily are the primary vectors of RNA viruses. These viruses include taxa with different RNA genomic structures and replication strategies, but all are non-retroviral viruses [3]. Collectively, these viruses are referred to as arthropod-borne (arbo-) viruses. Within the *Aedes* genus, *Aedes aegypti* and *Aedes albopictus* are the main arboviral vectors due to their broad geographic distribution, adaptation to breed in human habitats, and the wide number of viral species from different genera that they can vector [4,5]. These two mosquito species are able to efficiently transmit arboviruses of the genera *Flavivirus* (e.g. dengue viruses [DENV], Zika virus [ZKV], Usutu, Japanese encephalitis and yellow fever viruses), *Alphavirus* (e.g. chikungunya virus [CHIKV]), viruses of the Venezuelan equine encephalitis [VEE] and eastern equine encephalitis [EEE] complexes), *Orthobunyavirus* (e.g. Potosi, Cache Valley and La Crosse virus [LACV]), *Phlebovirus* (e.g. Rift Valley fever virus [RVFV]) and *Orbivirus* (e.g. Orungo Virus) [5-7]. Mosquitoes of the *Culex pipiens* complex, such as *Cx. pipens pipiens* and *Cx. quinquefasciatus*, are the most prominent *Culex* vectors because of their wide distribution and close association with humans [7]˙.These mosquito species are primary vectors of encephalitic flaviviruses, such as West Nile virus (WNV) and Japanese encephalitis virus, and they can also vector RVFV [7,8]. The only arbovirus know to be transmitted by Anophelinae is the alphavirus O’nyong-nyong [9]. Recently, additional RNA viruses have been identified from wild mosquitoes, but their virulence to humans and their impact on vector competence is still uncertain [9-11].

Mosquito competence for arboviruses is a complex and evolving phenotype because it depends on the interaction of genetic factors from both mutation-prone RNA viruses and mosquito vectors with environmental variables [12-15]. Not surprisingly, large variation exists in vector competence not only among mosquito species, but also across geographic populations within a species [16,17]. Understanding the genetic components of vector competence and how these genetic elements are distributed in natural populations and interact with environmental factors is essential for predicting the risk of arboviral diseases and for developing new transmission-blocking strategies [12]. Genomic and functional studies, primarily in *Drosophila melanogaster* and *Aedes* mosquitoes, have shown that RNA interference (RNAi) is the main antiviral mechanism in insects [18-20]. In this pathway, small RNAs are used to guide a protein-effector complex to target RNA based on sequence-complementarity. Three RNA silencing mechanisms exist: the microRNA, small interfering RNA (siRNA) and PIWI-interacting RNA (piRNA) pathways. They can be distinguished based on the mechanism of small RNA biogenesis and the effector protein complex to which these small RNAs associate [18,19]. While the role of the siRNA pathway in restricting arboviral infection has been widely studied and appears universal across mosquitoes, recent studies highlight the contribution of the piRNA pathway in antiviral immunity of *Aedes* mosquitoes [21]. Although important aspects of piRNA biogenesis and function in mosquitoes remains to be elucidated, it is clear that endogenous piRNAs arise from specific genomic loci called piRNA clusters, as was originally observed in *D. melanogaster* [22]. These piRNA clusters contain repetitive sequences, remnants of transposable elements and, in *Ae. aegypti*, virus-derived sequences [23].

Recent studies have shown that the genomes of some eukaryotic species, including mosquitoes, carry integrations from non-retroviral RNA viruses [24-32]. Viral integrations are generally referred to as Endogenous Viral Elements (EVEs) [33] or, if they derive from non-retroviral RNA viruses, as Non-Retroviral Integrated RNA Viruses Sequences (NIRVS) [29,34]. Integration of non-retroviral sequences into host genomes is considered a rare event because it requires reverse transcription by an endogenous reverse transcriptase, nuclear import and genomic insertion of virus-derived DNA (vDNA) [35]. During infection with DENV, WNV, Sindbis virus, CHIKV and LACV, fragments of RNA virus genomes are converted into vDNA by the reverse transcriptase activity of endogenous transposable elements (TEs) in cell lines derived from *D. melanogaster, Culex tarsalis, Ae. aegypti*, and *Ae. albopictus*, as well as in adult mosquitoes. The episomal vDNA forms produced by this mechanism reside in the nucleus and it has been proposed that they contribute to the establishment of persistent infections through the RNAi machinery [20,36-37]. These recent studies not only show that reverse transcription of RNA viruses occurs in Culicinae, they also suggest the functional involvement of RNAi.

Here we used a bioinformatics approach to analyze the presence, abundance, distribution, and transcriptional activity of NIRVS across the currently available 22 mosquito genome sequences. We probed these genomes for integrations from 425 non-retroviral viruses, including 133 arboviruses. We observed a ten-fold difference in the number of NIRVS between *Aedes* and the other tested mosquitoes. NIRVS were not evenly distributed across *Aedes* genomes, but occurred preferentially in piRNA clusters and, accordingly, they produced piRNAs. Among the viral species tested, integrations had the highest similarities to rhabdoviruses, flaviviruses and bunyaviruses, viruses that share the same evolutionary origin [38].

The larger number of NIRVS identified in *Ae. aegypti* and *Ae. albopictus*, their genome locations and their production of piRNAs show that in these species genomic integrations of viral sequences is a more pervasive process than previously thought and we propose that viral integrations contributes to shape vector competence.

## RESULTS

### NIRVS are unevenly distributed across mosquito species

25 genome assemblies currently available for 22 Culicinae species, along with the genome of *D. melanogaster*, were searched bioinformatically for sequence integrations derived from all 424 non-retroviral RNA viruses for which a complete genome sequence is currently available. Additionally, we tested the genome of African Swine Fever Virus, the only known DNA arbovirus [3], giving a total of 133 arboviruses (Figure 1, Additional files 1 and 2: Supplemental Tables S1 and S2). The genomes of 16 individual *Ae. albopictus* Foshan mosquitoes were sequenced to further validate NIRVS in this species. Retrieved sequences longer than 100 base pairs (bp) were filtered based on gene ontology and the presence of partial or complete open reading frames (ORFs) of viral proteins. This stringent pipeline led to the characterization of a total of 242 loci harboring NIRVS across the genome of 15 mosquitoes (Table 1, Figure 2). NIRVS loci were unevenly distributed across species. Anopheline species had a maximum of 7 NIRVS-loci, one NIRVS-locus was found in *Cx. quinquefasciatus*, 122 NIRVS were detected in *Ae. aegypti*, and 72 were found in *Ae. albopictus*. The NIRVS landscape was highly variable across the 16 *Ae. albopictus* sequenced genomes with extensive differences in the number of NIRVS and in their length, suggesting that NIRVS are frequently rearranged (Figure 3). No read coverage was observed in any of the 16 sequenced genomes for a total of 10 integrations that had been identified bioinformatically from the genome assembly of the Foshan strain (Additional file 3: Supplemental Table S3). The percentage of mapped reads and coverage was comparable across libraries excluding insufficient sequence depth as an explanation for the differential presence of NIRVS (Additional file 4: Supplemental Table S4). It is currently unclear if these 10 NIRVS are rare integrations or result from mis-assembly of the reference genome. Among the 11 viral families tested, NIRVS had sequence similarities exclusively with viruses of the *Rhabdoviridae, Flaviviridae, Bunyaviridae* and *Reoviridae* families, including currently circulating viruses (Table 1). Reoviridae- and Bunyaviridae-like integrations were similar to recently characterized viruses [39,40] and were rare, with no more than one integration per species (Figure 2). Phylogenetic analyses showed that viral integrations from Reoviridae were separated from currently known viral species in this family (Figure 4A,B). Integrations from *Bunyaviridae* were at the base of the phylogenetic tree and clustered with newly identified viruses such as Imjin virus and Wutai mosquito virus [40,41] (Figure 4C). In contrast, we observed numerous integrations from viruses of different genera within the Rhabdoviridae family and from viruses of the *Flavivirus* genus in multiple mosquito species, predominantly in *Ae. aegypti* and *Ae. albopictus* (Figure 2). *Rhabdoviridae*-like NIRVS (R-NIRVS) had similarities to genes encoding Nucleoprotein (N), Glycoprotein (G) and the RNA-dependent RNA polymerase (L), the relative abundance of which differed across mosquito species. We did not detect integrations corresponding to the matrix (M) or phosphoprotein (P) genes, consistent with observations in other arthropods [28]. R-NIRVS from Culicinae and Anophelinae formed separate clades in phylogenetic trees, supporting the conclusion that independent integrations occurred in the two mosquito lineages (Figure 4E). *Flavivirus*-like NIRVS (F-NIRVS) with similarities to structural genes (envelope [E], membrane [prM] and capsid [C]) were less frequent than integrations corresponding to non-structural genes (Figure 2). Some R-NIRVS or F-NIRVS sequences within one mosquito genome were highly similar to each other (nucleotide identity > 90%, (Additional file 5: Supplemental Table S5), which suggest that these were duplicated in the genome after a single integration event. This interpretation is also supported by the genomic proximity of several of these NIRVS (Figure 5). Surprisingly, identical NIRVS in *Ae. aegypti* were found not only adjacent to one another, but also at locations that are physically unlinked (i.e. AeRha138, AeRha110 and AeRha111). Thus, we cannot determine whether these identical NIRVS represent recent independent integration events or arose from duplication or ectopic recombination after integration.

**Figure 1.**
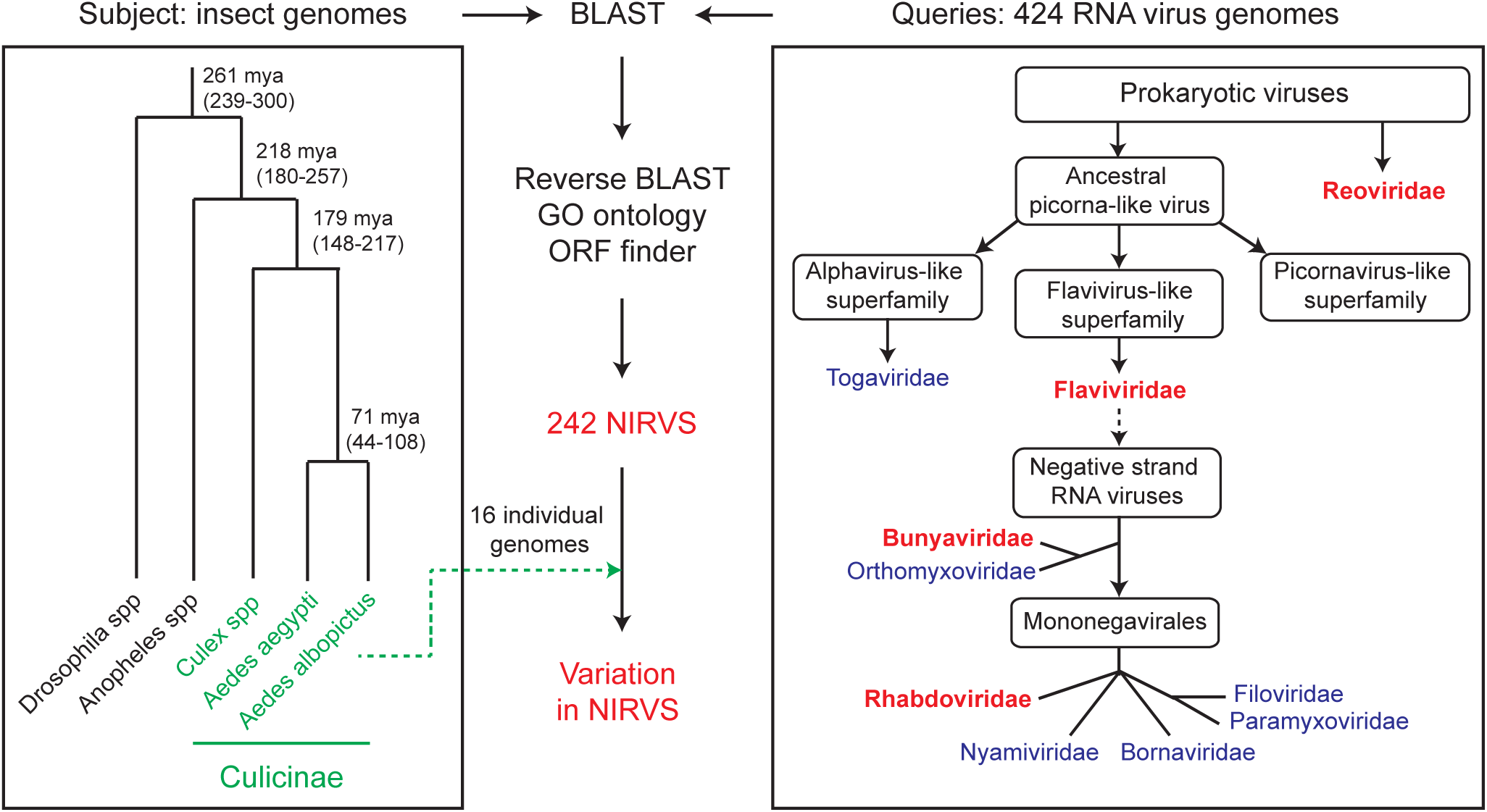
Pipeline for NIRVS identification. The currently available 22 mosquito genomes and the genome of *Drosophila melanogaster* were probed bioinformatically using tblastx and 425 viral species (424 non-retroviral RNA viruses and 1 DNA arbovirus). Tested insect and viral RNA genomes are shown in the context of their phylogeny [2,38]. Identified blast hits were parsed based on gene ontology and the presence of partial or complete viral ORFs. In *Ae. albopictus*, bioinformatic analyses was extended to whole-genome sequencing data from 16 individual mosquitoes of the Foshan strain. This stringent pipeline led to the characterization of 242 loci with NIRVS. Viral families for which NIRVS were characterized are shown in red.

**Figure 2.**
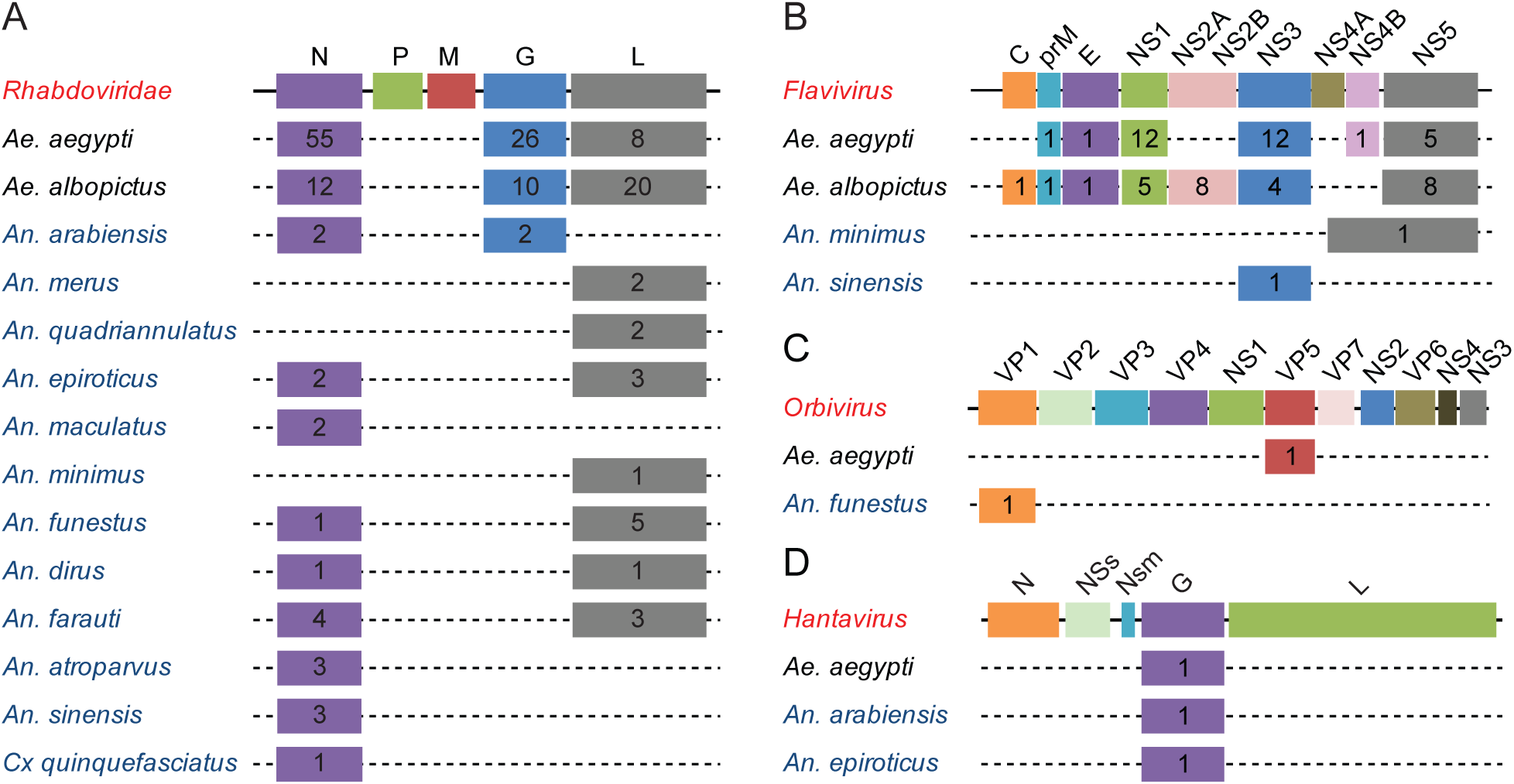
Different abundance of NIRVS across virus genera, genes and host species. Schematic representation of the genome structures of *Rhabdoviridae* (A) and the genera *Flavivirus* (family *Flaviviridae*) (B), *Orbivirus* (family *Reoviridae*) (C) and *Hantavirus* (Family *Bunyaviridae*) (D). Numbers within each box represent the number of NIRVS loci spanning the corresponding viral gene per mosquito species. When a NIRVS locus encompassed more than one viral gene, the viral gene with the longest support was considered. Mosquitoes of the Culicinae and Anophelinae subfamilies are in black and blue, respectively. Dotted lines indicate viral integrations were not contiguous in the host genomes.

**Figure 3.**
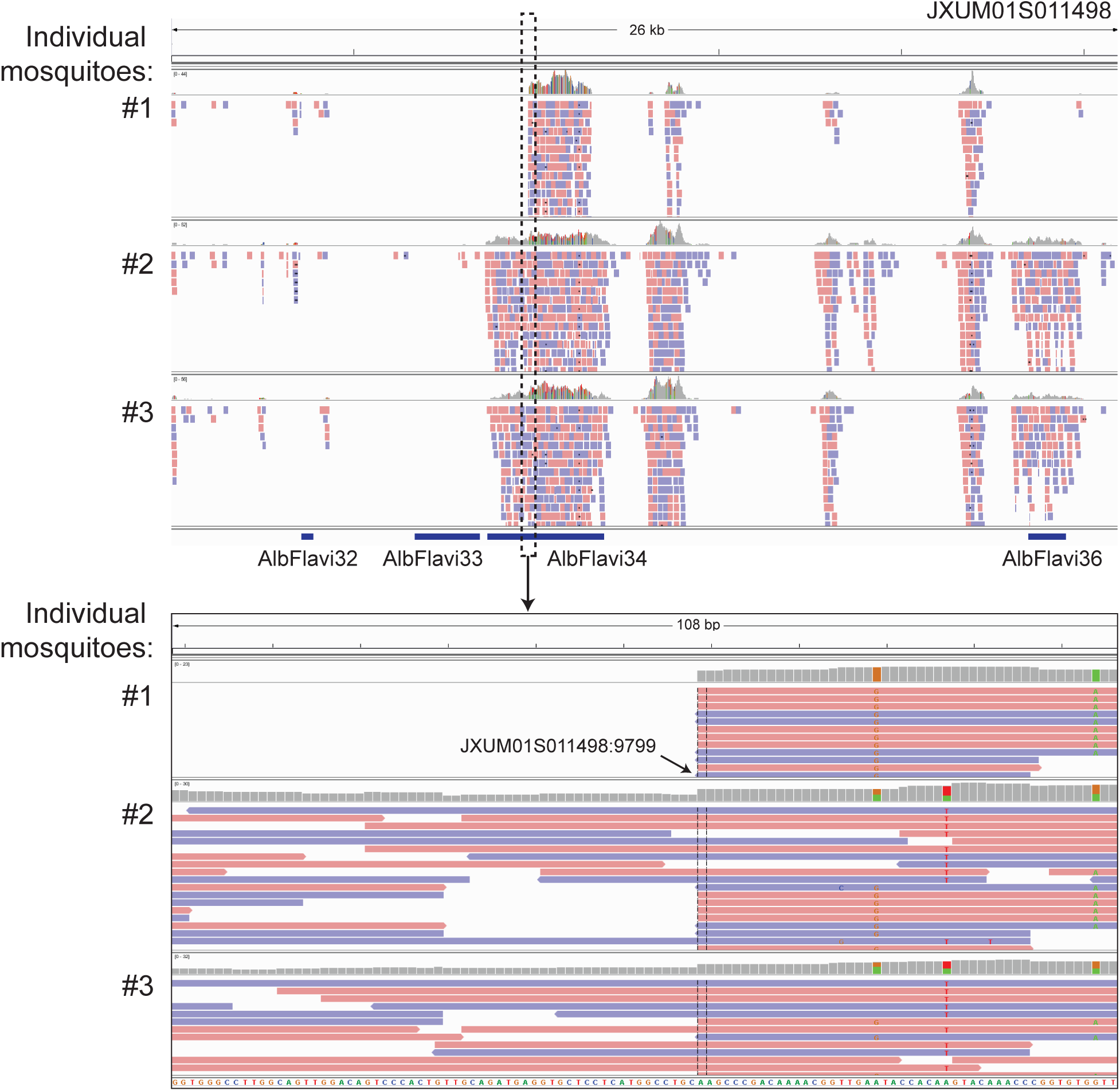
Variability of NIRVS within the *Ae. albopictus* Foshan strain. Bioinformatic analyses of the *Ae. albopictus* genome identified 4 NIRVS on scaffold JXUM01S011498: AlbFlavi32, AlbFlavi33, AlbFlavi34 and AlbFlavi36. No read coverage was seen for AlbFlavi32 and AlbFlavi33 in any of the 16 sequenced genomes. AlbFlavi36 had read coverage in 13 of the 16 tested mosquitoes, whereas AlbFlavi34 showed length variability.

**Figure 4.**
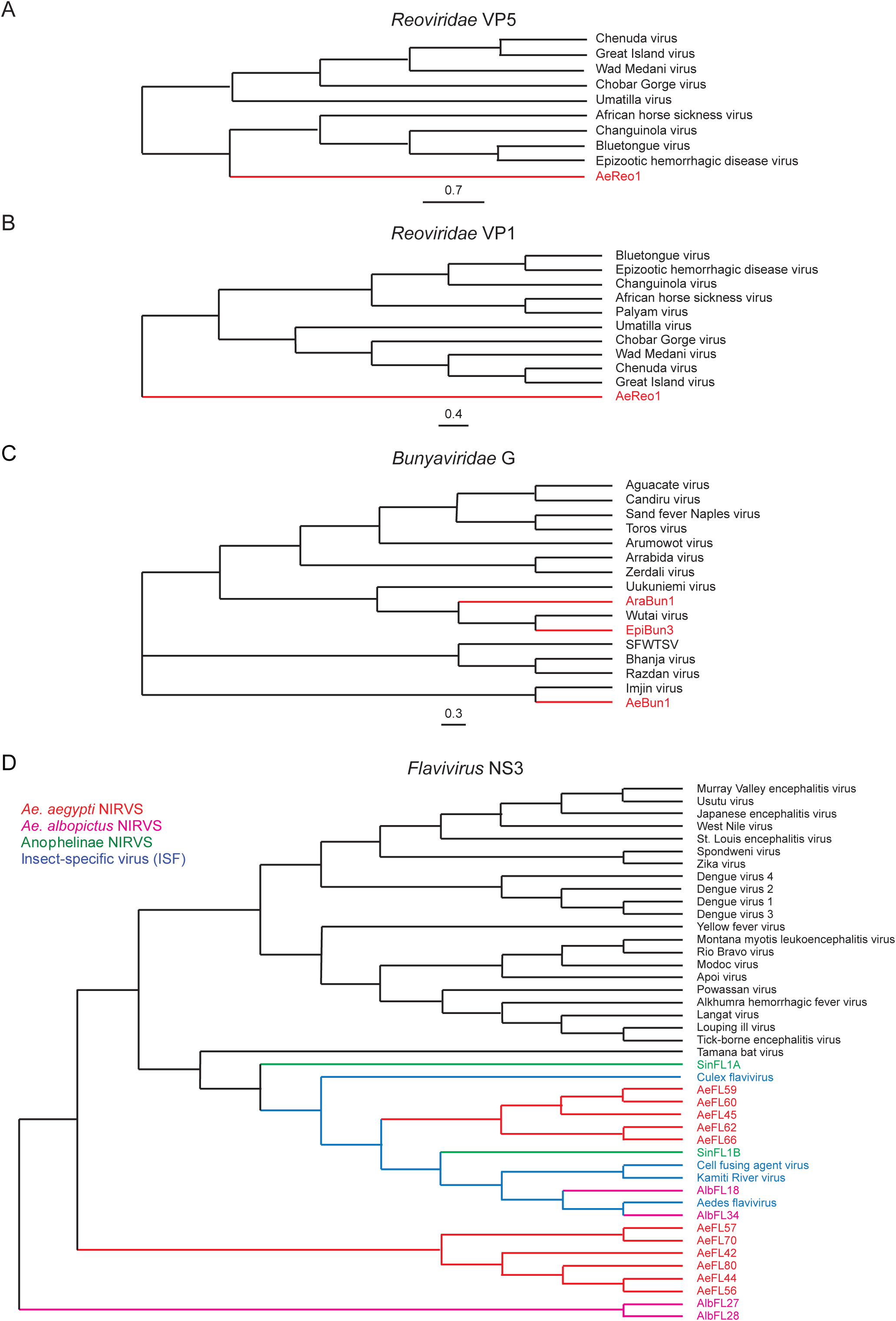
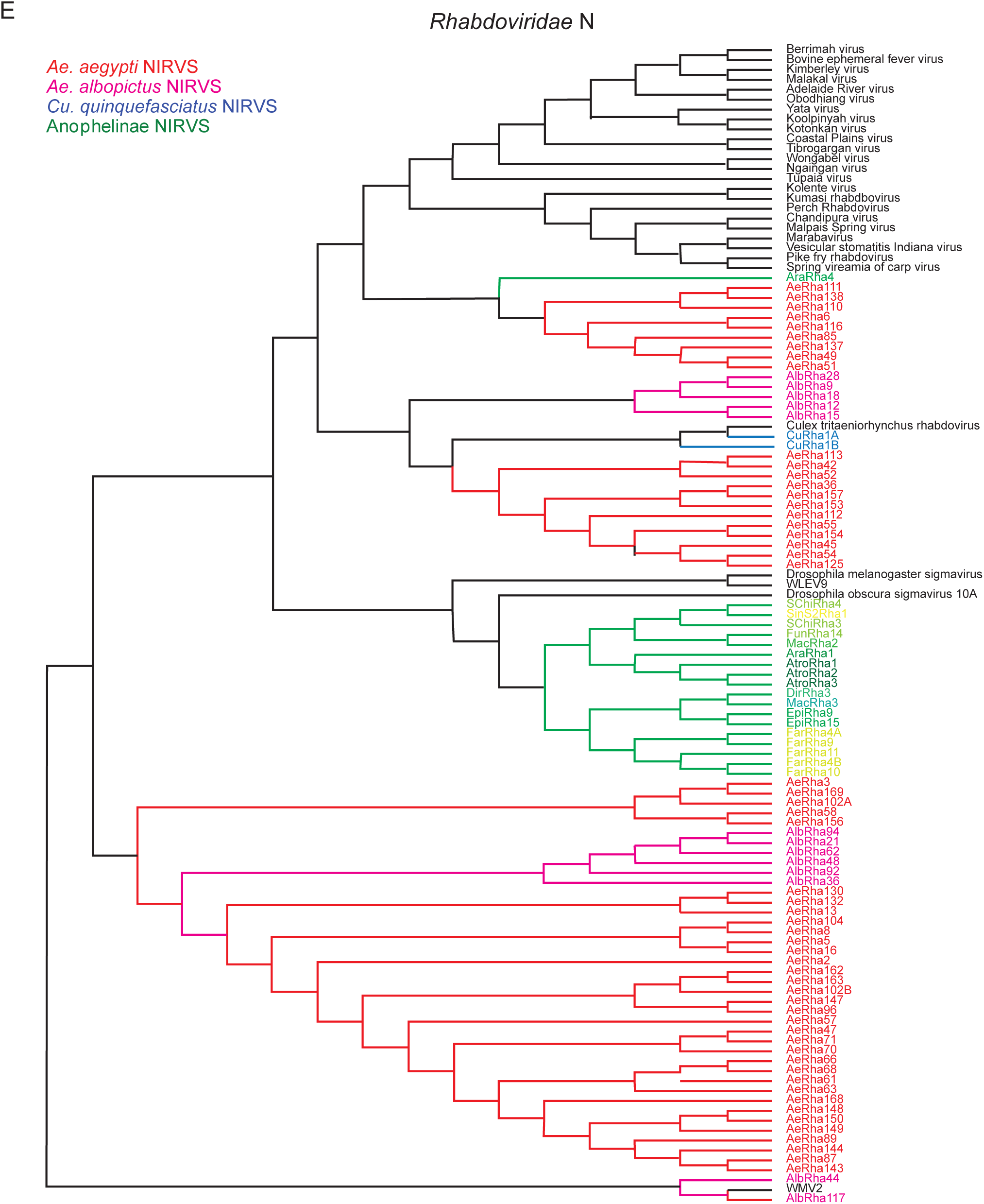
Phylogenetic analyses of *Reoviridae, Bunyaviridae, Flavivirus*, and *Rhabdoviridae-*like integrations. Phylogenetic relationships of NIRVS with similarity to the *Reoviridae* VP5 (A), *Reoviridae* (B), *Bunyaviridae* G (C),*Flavivirus* NS3 (D), and *Rhabdoviridae* N (E) genes. The evolutionary history was inferred using the Maximum Likelihood method. The trees with the highest log likelihood are shown. Support for tree nodes was established after 1000 bootstraps.

**Figure 5.**
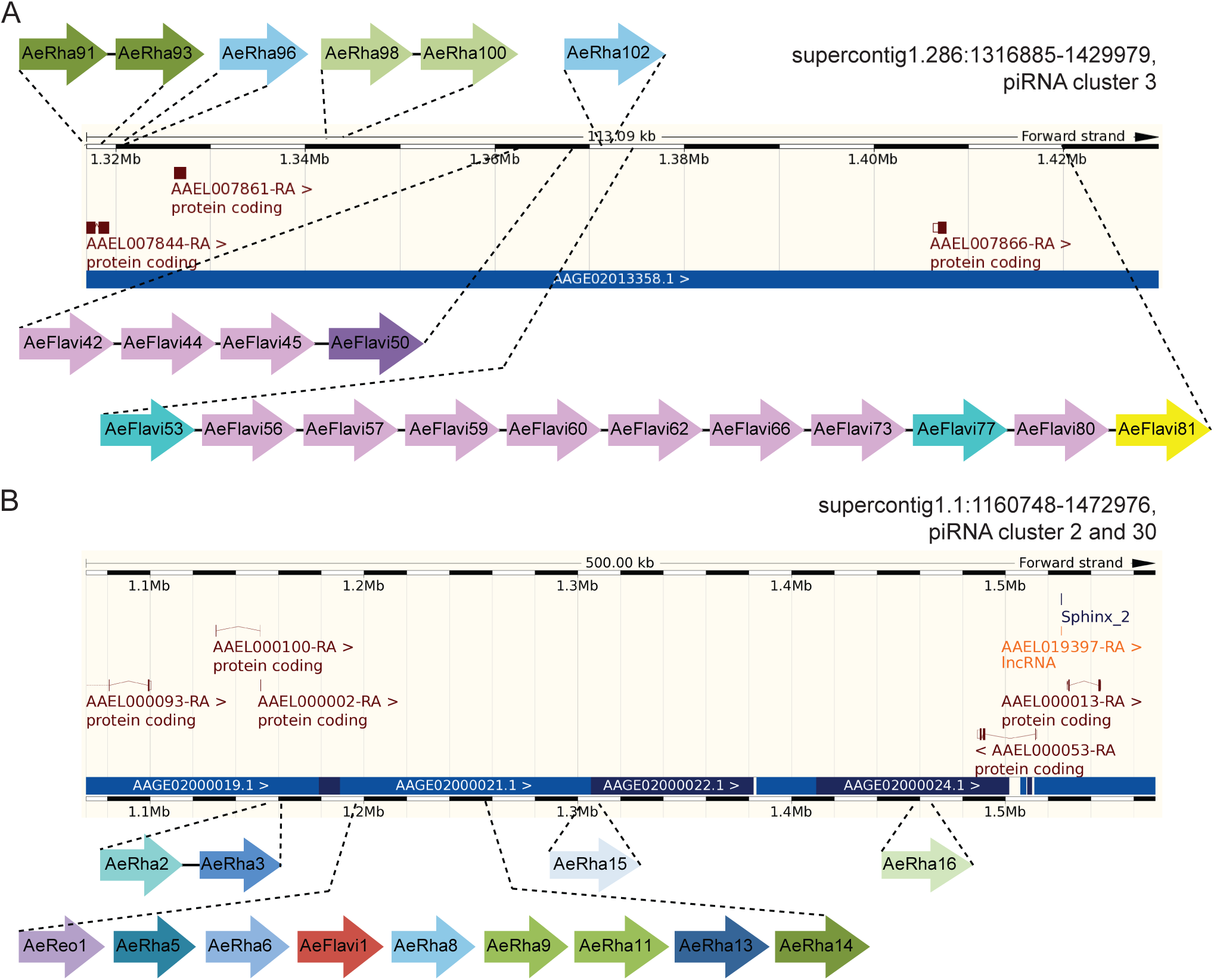
Enrichment of NIRVS in two regions of the *Ae. aegypti* genome. One fourth of the identified NIRVS in *Ae. aegypti* map to two genomic regions. (A) Region 1 (supercont1.286:1316885-1429979) includes piRNA cluster 3 [23] and is enriched in the LTR transposons LTR/Pao_Bel and LTR/Ty3_gypsy, which occupy 16.33 and 14.98% of the region, respectively. (B) Region 2 (supercont1.1:1160748-1472976) includes piRNA clusters 2 and 30 and is also enriched for LTR transposons. LTR/Ty3_gypsy occupancy in region 2 is 24.18%. NIRVS are color-coded based on their sequence identity (Additional file 5: Supplemental Table S5).

**Table 1.** Number of viral integrations (NIRVS) detected for each of the viral families (N. species) tested across the 22 mosquito genomes. A total of 424 non-retroviral RNA viruses with complete genomes were analyzed. The genome of African swine fever virus, the only known DNA arbovirus was also included in the analyses, but no NIRVS were found for this virus. Mosquito species and viral families for which NIRVS were detected are in bold.

| Mosquito species | Families of tested non-retroviral RNA viruses<sup>†</sup> (N. species) |  |  |  |  |  |  |  |  |  |
| --- | --- | --- | --- | --- | --- | --- | --- | --- | --- | --- |
|  | T*<br>(24) | F*<br>(92) | B*<br>(59) | R*<br>(70) | O*<br>(4) | Rha*<br>(93) | Bo<br>(6) | Fi<br>(8) | N<br>(4) | P<br>(64) |
| *Aedes aegypti* | 0 | **32** | **1** | **1** | 0 | **88** | 0 | 0 | 0 | 0 |
| *Aedes albopictus* | 0 | **30** | 0 | 0 | 0 | **42** | 0 | 0 | 0 | 0 |
| *Culex quinquefasciatus* | 0 | 0 | 0 | 0 | 0 | **1** | 0 | 0 | 0 | 0 |
| *Anopheles christy* | 0 | 0 | 0 | 0 | 0 | 0 | 0 | 0 | 0 | 0 |
| *Anopheles gambiae* | 0 | 0 | 0 | 0 | 0 | 0 | 0 | 0 | 0 | 0 |
| *Anopheles coluzzi* | 0 | 0 | 0 | 0 | 0 | 0 | 0 | 0 | 0 | 0 |
| *Anopheles arabiensis* | 0 | 0 | **1** | 0 | 0 | **4** | 0 | 0 | 0 | 0 |
| *Anopheles melas* | 0 | 0 | 0 | 0 | 0 | 0 | 0 | 0 | 0 | 0 |
| *Anopheles merus* | 0 | 0 | 0 | 0 | 0 | **2** | 0 | 0 | 0 | 0 |
| *Anopheles quadrianulatus* | 0 | 0 | 0 | 0 | 0 | **2** | 0 | 0 | 0 | 0 |
| *Anopheles epiroticus* | 0 | 0 | 0 | 0 | 0 | **7** | 0 | 0 | 0 | 0 |
| *Anopheles stephensi* | 0 | 0 | 0 | 0 | 0 | 0 | 0 | 0 | 0 | 0 |
| *Anopheles maculatus* | 0 | 0 | 0 | 0 | 0 | **2** | 0 | 0 | 0 | 0 |
| *Anopheles culicifacies* | 0 | 0 | 0 | 0 | 0 | 0 | 0 | 0 | 0 | 0 |
| *Anopheles minimus* | 0 | **1** | 0 | 0 | 0 | **1** | 0 | 0 | 0 | 0 |
| *Anopheles funestus* | 0 | 0 | 0 | **1** | 0 | **7** | 0 | 0 | 0 | 0 |
| *Anopheles dirus* | 0 | 0 | 0 | 0 | 0 | **4** | 0 | 0 | 0 | 0 |
| *Anopheles farauti* | 0 | 0 | 0 | 0 | 0 | **7** | 0 | 0 | 0 | 0 |
| *Anopheles atroparvus* | 0 | 0 | 0 | 0 | 0 | **3** | 0 | 0 | 0 | 0 |
| *Anopheles sinensis* | 0 | **1** | 0 | 0 | 0 | **1** | 0 | 0 | 0 | 0 |
| *Anopheles albimanus* | 0 | 0 | 0 | 0 | 0 | 0 | 0 | 0 | 0 | 0 |
| *Anopheles darlingi* | 0 | 0 | 0 | 0 | 0 | 0 | 0 | 0 | 0 | 0 |

T=Togaviridae; F=Flaviviridae; B=Bunyaviridae; R=Reoviridae; O=Orthomyxoviridae; Rha=Rhabdoviridae; Bo=Bornaviridae; Fi=Filoviridae; N=Nyamiviridae; P=Paramyxoviridae. \*viral families that contain arboviruses

Generally, NIRVS were most similar to insect-specific viruses (ISVs), which replicate exclusively in arthropods, but are phylogenetically-related to arboviruses [10,42] (Figure 4D). However, we observed integrations that were most similar to arboviruses of the *Vesiculovirus* genus (*Rhabdoviridae*) in both *Ae. aegypti* and *Ae. albopictus* (Additional File: Supplemental Table S3).

### NIRVS produce piRNAs and map in piRNA clusters more frequently than expected by chance

To better understand the mechanisms of integration, we analyzed in greater detail the genomic context of NIRVS in *Ae. aegypti* and *Ae. albopictus*, the mosquitoes with the largest number of identified NIRVS. Previously, uncharacterized viral sequences were identified as piRNA producing loci in *Ae. aegypti* [23,43], and these observations prompted us to analyze whether NIRVS are enriched in piRNA clusters. Currently annotated piRNA clusters represent 1.24% and 0.61% of the *Ae. aegypti* and *Ae. albopictus* genomes, respectively [23,44]. Remarkably, 44% and 12.5% of all NIRVS map to these genomic loci, and these frequencies are significantly higher than expected by chance (Table 2). Enrichment of NIRVS in piRNA clusters in *Ae. aegypti* was driven by two regions that harbored one fourth of all NIRVS loci (region1: scaffold 1.286: 1316885-1429979; region 2: scaffold 1.1: 1160748-1472976), which includes piRNA cluster 3 and piRNA clusters 2 and 30, respectively [23]. In these two regions, NIRVS span partial ORFs with similarities to different *Rhabdovirus* and *Flavivirus* genes, with instances of duplications as well as unique viral integrations (Figure 5). NIRVS also were enriched in regions annotated as exons in *Ae. albopictus*, but not in *Ae. aegypti* (Table 2).

**Table 2.** Clustering of viral integrations (NIRVS) in piRNA loci of the *Ae. aegypti* (A) and *Ae. albopictus* (B)genomes. The probability (P) of observing *k* NIRVS loci in piRNA clusters, coding genes and intergenic regions. P was estimated using cumulative binomial distribution; a value of P < 0.05 indicates a statistically significant enrichment of NIRVS in the corresponding genomic region.

| Host | Genomic region | Length (bp) | % genome | k integrations* | P |
| --- | --- | --- | --- | --- | --- |
| *Ae. aegypti* | piRNA cluster | 17,000,000 | 1.24 | 54 | $< 10^{-10}$ |
|  | Coding genes | 286,538,182 | 20.82 | 24 | 0.66 |
|  | Intergenic regions | 1,072,461,818 | 77.94 | 44 | 1 |
| *Ae. albopictus* | piRNA cluster | 1,926,670 | 0.61 | 9 | $< 10^{-10}$ |
| | Coding genes | 163,407,667 | 8.26 | 14 | $2.08 \times 10^{-3}$ |
|  | Intergenic regions | 1,803,592,333 | 91.14 | 49 | 1 |

\*6 integrations in the *Ae. aegypti* genome were in exons of genes within piRNA clusters; in this analyses they were attributed to piRNA clusters. Statistical significance did not change when these integrations were assigned to coding genes

The presence of NIRVS in piRNA clusters prompted us to analyze the expression of NIRVS-derived small RNAs. Therefore we used deep-sequencing data from published resources and mapped small RNAs on NIRVS sequences after collapsing those elements that shared identical sequences (Additional File 5: Table S5). Small RNAs in the size range of piRNAs (25-30 nucleotides), but not siRNAs (21-nucleotides) mapped to NIRVS in both *Ae. aegypti* and *Ae. albopictus*, independently of genomic localization and corresponding viral ORFs, (Figure 6A,B). Generally, piRNAs derived from individual NIRVS sequences are not highly abundant. Of all tested NIRVS, 43% (n=33) and 11% (n=6) had at least 10 piRNA reads per million genome-mapped reads in *Ae. aegypti* and *Ae. albopictus*, respectively. In *Ae. aegypti*, the highest piRNA counts were a few hundred reads per million genome-mapped reads. In *Ae. albopictus* the maximum piRNA counts per NIRVS were about 10 fold lower, suggesting that NIRVS piRNA are less efficiently produced or retained in this species. In both mosquito species, R-NIRVS showed higher coverage than F-NIRVS (Figure 6E). These piRNAs were biased for uridine at position 1 and primarily in antisense orientation to the predicted viral ORF, establishing the potential to target viral mRNA (Figure 6A-D). Yet, a 10A bias of sense piRNAs, particularly in *Ae. albopictus* indicates some NIRVS produce piRNAs through ping-pong amplification. Interestingly, ping-pong dependent secondary piRNAs seem to be almost exclusively (100% in *Ae. aegypti* and >99.5% in *Ae. albopictus*) derived from R-NIRVS (Figure 6E). The nature of this specific induction of secondary piRNAs biogenesis from Rhabdoviral sequences is currently unknown.

**Figure 6.**
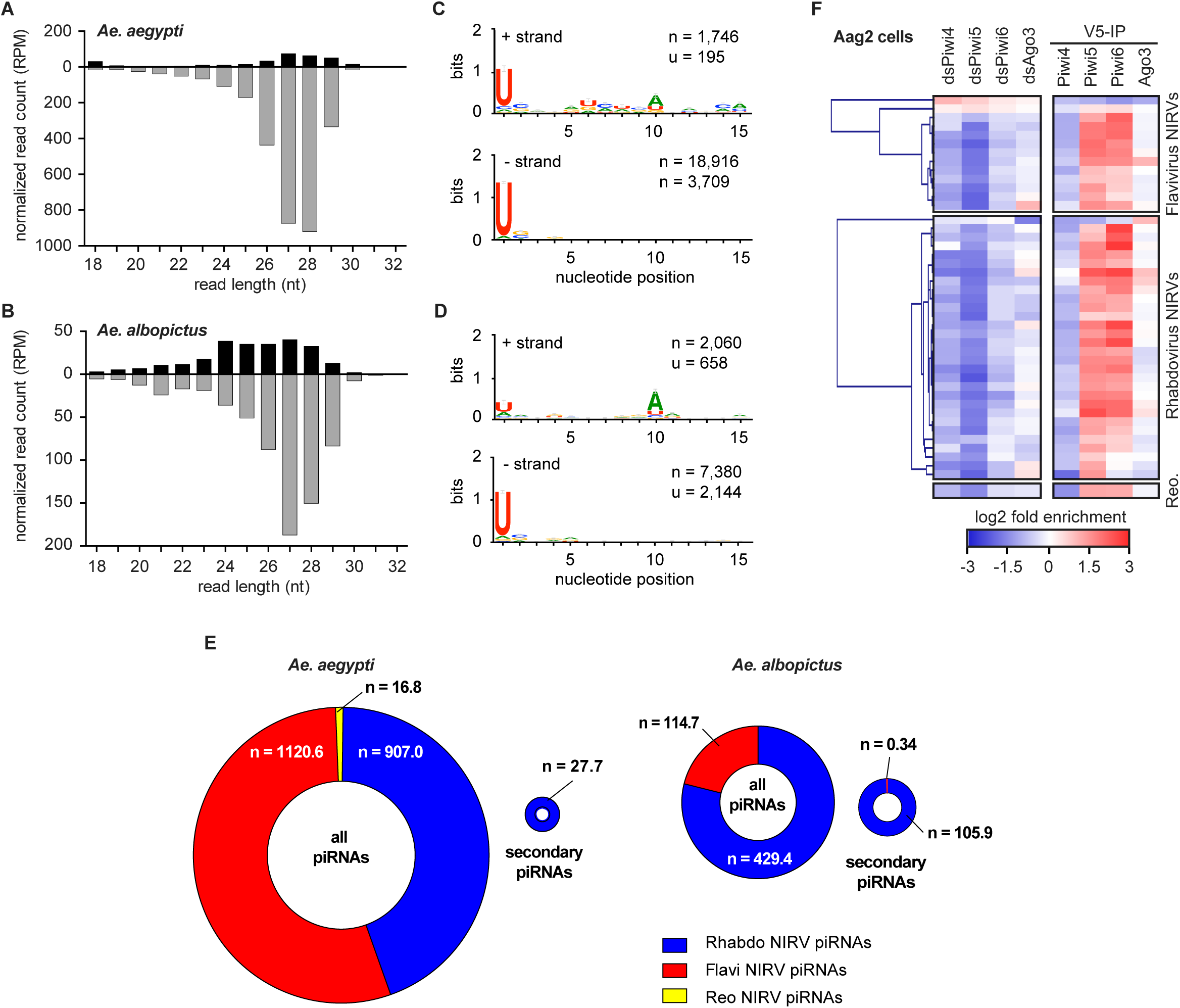
NIRVS produce 25-30 nt piRNAs, but not 21-nt siRNAs. Size distribution of small RNAs from published resources mapping to NIRVS in the *Ae. aegypti* (A) and *Ae. albopictus* (B) genomes. NIRVS-derived piRNAs are biased for sequences that are antisense to viral mRNAs, suggesting potential to target viral RNA. NIRVS-derived piRNAs are biased for uridine at position 1, in both *Ae. aegypti* (C) and *Ae. albopictus* (D). (E) Number of all piRNAs and secondary piRNAs expressed in *Ae. aegypti* or *Ae albopictus*. Ring charts are drawn to scale and numbers indicate the normalized piRNA counts of F-NIRVS (red), R-NIRVS (blue), and NIRVS from *Reovirus* that have been found only in *Ae. aegypti* (yellow). (F) Left panel, heat map of the relative abundance ofNIRVS-derived small RNAs in Aag2 cells in which PIWI expression was silencing using RNAi (dsPiwi4-6, and dsAgo3), compared to control dsRNA treatment. Right panel, heatmap of small RNA enrichment in immunoprecipitations (IP) in the indicated PIWI proteins over control GFP IP.

We next analyzed the dependency on and association with PIWI proteins of NIRVS-derived small RNAs in Aag2 cells [45] and found that small RNA expression was reduced by knockdown of Piwi5 and, to a lesser extent, Piwi4 and Piwi6 (Figure 6F), with only few exceptions. Consistent with this finding, NIRVS-derived small RNAs were most enriched in immunoprecipitations (IP) of Piwi5 and Piwi6 (Figure 6F). Together, these data indicate that NIRVS produce piRNAs, the majority of which have the characteristics of primary piRNAs. Yet, secondary piRNA biogenesis as indicated by a 10A bias and association with Ago3, seems to occur specifically from R-NIRVS.

### NIRVS and transposable elements

piRNA clusters in *Dr. melanogaster* are enriched for remnants of TE sequences, and it is likely that vDNA is produced by the reverse transcriptase activity of TEs [36,37]. Moreover, NIRVS-derived piRNAs resembled the characteristics of TE-derived piRNAs in their antisense 1U bias and enrichment in Piwi5 and Piwi6 protein complexes. We analyzed the transposon landscape of NIRVS loci by systematically identifying all annotated TEs in the 5 and 10 kb genomic regions flanking each side of the NIRVS integration. We observed that NIRVS were predominantly associated with long terminal repeat (LTR) retrotransposons. Within LTR-retrotransposons, we observed enrichment of members of the Ty3_gypsy families (Table 3). Such enrichments were even more pronounced in the two regions in *Ae. aegypti* where 40% of NIRVS reside (Figure 5). While LTR retrotransposon occupancy was 12.34% across the entire *Ae. aegypti* genome, it reached 23.60-25.88%, 31.35%, and 30.55% in regions flanking all NIRVS-loci, region 1, and region 2, respectively. More strikingly, while the Ty3_gypsy families of LTR retrotransposon occupancy was 2.58% across the entire *Ae. aegypti* genome, it reached 14.7-17.5%, 14.98% and 24.18% in regions flanking all NIRVS-loci, region 1, and region 2, respectively (Table 3). Nine full-length TEs were found flanking NIRVS-loci, seven of which are Ty3_gypsy retrotransposons. For example, 3 copies and 1 copy of the full-length Ty_gypsy_Ele58 (TF000321) were found in regions 1 and 2, respectively. Moreover, one viral integration in *Ae. aegypti* (i.e. AeBunya1) was found embedded within a full-length TE of the Pao-Bel family.

**Table 3.** NIRVS and transposable elements (TEs)Es. Analyses of TE enrichment througout the *Ae. aegypti* genome (AaegL3), in regions harboring NIRVS (NIRVS), in region1 and in region 2, respectively.

| TE group<sup>1</sup> | TE Occupancy (%) |  |  |  |
| --- | --- | --- | --- | --- |
|  | AaegL3 | NIRVS<sup>3</sup> | Region 1<sup>4</sup> | Region 2<sup>5</sup> |
| LTR retrotransposons | 12.34 | 23.06 (25.88) | 31.35 | 30.56 |
| LTR/Pao_Bel | 4.42 | 6.9 (6.49) | 16.33 | 4.15 |
| LTR/Ty1_copia | 5.34 | 1.46 (1.90) | 0.04 | 2.22 |
| LTR/Ty3_gypsy | 2.58 | 14.7 (17.50) | 14.98 | 24.18 |
| non-LTR retrotransposons | 12.81 | 3.91 (5.54) | 0 | 3.43 |
| SINEs | 1.14 | 0.16 (0.22) | 0 | 0 |
| DNA transposons | 6.96 | 3.29 (3.49) | 3.38 | 2.72 |
| MITEs | 12.81 | 8.03 (7.34) | 0.26 | 2.34 |
| Helitrons | 1.2 | 2.01 (2.32) | 0 | 2.04 |
| Penelope | 0.42 | 0.2 (0.28) | 0.26 | 0.8 |

<sup>1</sup> For consistency with previous publication and for clear classification, only TEs annotated in TEfam are used. We used TE occupancy (number of bases in the genomic sequence that match TEs) as an indication for possible TE enrichment. TE copy number was not used because to prevent that TEs that are broken into multiple fragments and counted multiple times.
<sup>2</sup> The genome assembly described in Nene et al. (2007) is slightly different from AaegL3 (*Aedes-aegypti*-Liverpool\_SCAFFOLDS\_AaegL3.fa), which is used in this analysis. For better comparison with viral integration sites, a new RepeatMasker analysis was performed using the AaegL3 assembly under the same default parameters.
<sup>3</sup> In addition to the integration site, 5 kb or 10 kb (in brackets) sequences flanking each side of the NIRVS were retrieved for the analysis. Because the NIRVS sequences are also included in the analyses, these results may be an under-estimate of the actual TE occupancy..
<sup>4</sup> TE analysis of viral integration sites (plus 5kb flanking each side) in Supercontig 1.286 between positions 1316885 bp and 1429979 bp.
<sup>5</sup> TE analysis of viral integration sites (plus 5kb flanking each side) in Supercontig 1.1 between positions 1160748 bp and 1472976 bp.

### NIRVS transcriptional activity

All NIRVS encompassed partial viral ORFs, with the exception of AlbFlavi34. AlbFlavi34 corresponds to a portion of the first *Flavivirus*-like sequence characterized in mosquitoes and includes a complete ORF for NS3 [24]. Two alleles of different lengths were seen for AlbFlavi34 in the 16-sequenced *Ae. albopictus* genomes (Figure 3). The short allele, which interrupts the NS3 ORF, had a frequency of 53% (Additional File 6: Supplemental file 1). Based on recent experimental data showing that NIRVS are transcriptionally active even if they do not encode a complete ORF [24,28,32,46] we analyzed NIRVS expression using published RNA-seq data from poly(A) selection protocols. Expression levels were < 5 reads per kilobase per million mapped reads (RPKM) for > 92% of all tested NIRVS, including NIRVS that produce piRNAs (Additional File 7: Table S6). Similar to small RNA profiles, expression levels of R-NIRVS were higher than those of F-NIRVS (Additional File 7: Table S6).

Despite RNA-seq data showing limited transcriptional activity for AlbFlavi34 (RPKM values ranging from 0.009 to 0.013), we analyzed its expression in different developmental stages by RT-qPCR using primers that amplify both the short and long alleles. Cycle threshold (Ct) values ranged from 27 (found in pupae) to 39.34 (detected in ovaries of blood fed-females) and 60% of the samples having Ct> 30, confirming low AlbFlavi34 expression. AlbFlavi34 expression was highest in the pupae and adult males in comparison to expression in larvae (Additional File 8: Figure S1). These data support the conclusions that steady-state RNA levels of most NIRVS are rather low or even undetectable. Yet, the production of piRNAs indicates that they must be transcriptionally active. Whether their precursor transcripts are non-polyadenylated or rapidly processed into piRNAs remains to be established.

## DISCUSSION

The genomes of mosquitoes and several eukaryotic species carry integrations from non-retroviral RNA viruses, including arboviruses. To shed light on the widespread and biological significance of this phenomenon, we analysed the presence, distribution and transcriptional activity of integrations from 424 non-retroviral RNA viruses, and one DNA arbovirus, in 22 mosquito genomes, in the context of both their phylogeny and mosquito vector competence. We showed that the arboviral vector species *Ae. aegypti* and *Ae. albopictus* have ten-fold more integrations than all other tested mosquitoes. Moreover, we found that viral integrations produce piRNAs and occur predominantly in piRNA clusters. Our results support the conclusion that the abundance of viral integrations is not dependent on viral exposure, but seems to correlate with the TE landscape and piRNA pathway of the mosquito.

### NIRVS viral origin

Across all 425 viral species tested, viral integrations had similarities primarily to ISVs of the *Bunyaviridae, Reoviridae* and, predominantly, *Rhabdoviridae* and *Flaviviridae* families. Notably, although the Togaviridae family contains mosquito-borne members as well as insect specific viruses, we identified no integrations from viruses of in this family. Further studies are required to clarify whether this result is due to a sampling bias or to the different evolutionary history of *Alphavirus*-like versus *Flavivirus-*like viruses [38]. For instance, Eilat virus and the Taï Forest alphavirus are the only insect-specific alphaviruses (family *Togaviridae*) identified and a large screen suggests that mosquito-specific viruses may not be abundant among alphaviruses [47] unlike the *Bunyaviridae, Reoviridae, Rhabdoviridae* and *Flaviviridae* families in which many ISVs have been identified [42]. An alternative explanation may be based on the interactions of these viruses with the piRNA machinery. For example, while both alphaviruses and flaviviruses produce vpiRNAs in *Aedes*, the distribution of piRNAs on the viral genomes are not comparable between these genera, suggesting that piRNA biogenesis might be different [21]. Both alphaviruses (Sindbis and CHIK viruses) and flaviviruses (DENV, WNV) have been shown to produce episomal vDNA forms that locate to the nucleus after infection of mosquitoes [20,36-37]. These vDNA forms do not arise uniformly from the whole viral genome and their profile may be different between alphaviruses and flaviviruses (Nag et al., 2016). If these episomal vDNA are integrated into the genome, a different vDNA profile will result in a different NIRVS landscape.

ISVs of the families *Bunyaviridae, Flaviviridae* and *Rhabdoviridae* families are ancient and diversified within their hosts, and they seem to be maintained in mosquitoes through transovarial transmission [10,42]. Additionally, mounting phylogenetic evidence implicate ISVs as precursors of arboviruses [48], for which vertical transmission occurs at a lower frequency than horizontal transmission through a vertebrate host [49]. Vertical transmission provides access to the mosquito germ-line, a mechanism through which NIRVS could be maintained within vector populations. Thus, the observed higher incidence of NIRVS from ISVs than arbovirus may be linked to differences in the frequency of their transovarial transmission.

NIRVS from *Bunyaviridae* and *Rhabdoviridae* have been identified in insects other than mosquitoes, including different *Drosophila* species and the tick *Ixodes scapularis* [26-28]. In contrast, NIRVS from *Flaviviruses* have been found only in mosquitoes, predominantly in *Ae. aegypti* and *Ae. albopictus* [2,26,32]. Interestingly, vertebrates that may be part of the arbovirus transmission cycle do not have integrations from arboviruses, but a low number (<10) of integrations from *Bornaviruses* and/or *Filoviruses* have been identified in humans, squirrel, microbat, opossum, lemur, wallaby and medaka [25-27]. Finally, several Anophelinae mosquitoes analyzed here were sampled in the same geographic area as *Ae. albopictus*, but showed 10 times fewer NIRVS than *Ae. albopictus* and *Ae. aegypti*. Overall, these data indicate that viral exposure is not a determinant of NIRVS, but that virus-host lineage-specific interactions play a crucial role in how their genomes co-evolve. Additionally, our comparative analysis shows that *Aedes* mosquitoes acquire and retain fragments of infecting non-retroviral RNA viruses primarily the *Flaviviridae* and *Rhabdoviridae* families, more frequently than other tested arthropods and vertebrates. A deeper understanding of the evolution of viruses within these large and diverse families, especially their recently characterized ISVs, along with insights into the variability of the genomes of mosquito populations are warranted to elucidate the dynamic species-specific interactions between RNA viruses and *Aedes* mosquitoes.

### NIRVS genomic context

NIRVS are significantly enriched in piRNA clusters in both *Ae. aegypti* and *Ae. albopictus*, which could be the result of positive selection favoring the retention of those NIRVS that integrated by chance in these genomic loci [50]. However, we also observed NIRVS in intergenic and coding sequences and found that NIRVS expressed piRNAs independently of their genomic localization. These observations suggest that additional piRNA clusters exist [23,44] or that other features in these NIRVS loci prime piRNAs production. For example, a piRNA trigger sequence (PTS) was recently found to drive piRNA production from a major piRNA cluster (named *Flamenco*) in *Drosophila* [51]. We analyzed the mosquito genome sequences, but we did not find PTS orthologous sequences in either *Ae. aegypti* nor *Ae. albopictus*. It remains to be established whether other PTS sequences exist that may explain piRNA production from non-cluster associated NIRVS.

Analyses of the integration sites showed that NIRVS are primarily associated with LTR transposons of the Gypsy and Pao families, which are the most abundant TE families in both *Ae. aegypti* and *Ae. albopictus* genomes [2]. Additionally, full-length TEs, primarily Ty3_gypsy retrotransposons, were are found to flank NIRVS-loci. This organization is compatible with recent experimental data showing that vDNA forms are produced by retrotransposon-derived reverse transcriptase, likely by template switching [20,37]. This arrangement also is favorable for ectopic recombination, a mechanism proposed for both NIRVS biogenesis and piRNA cluster evolution [52]. Ectopic recombination would be a more parsimonious explanation than independent integrations from the same viral source for our finding of several not physically-linked, but identical *Ae. aegypti* NIRVS. Despite many remaining uncertainties due to the highly repetitive and complex structure of the regions in which NIRVS map, these data confirm a functional link among NIRVS, TEs, and the piRNA pathway.

### NIRVS and mosquito immunity

Our data indicate that in *Ae. aegypti* and *Ae. albopictus* NIRVS do not encode proteins that interfere *in trans* with viral products as was observed in bornavirus-derived NIRVS in vertebrates [53]. Rather our data suggest that NIRVS may be part of a piRNA-based antiviral response. Only one of the characterized NIRVS had a complete viral ORF, which showed two alleles of different length within the 16 individuals of the Foshan strain that we sequenced. The short variant interrupted the NS3 ORF. We cannot exclude that this is due to lack of purifying selective pressure as the *Ae. albopictus* Foshan strain has been reared under standard laboratory conditions without infection challenges for more than 30 years [2]. However, the enrichment of NIRVS within piRNA clusters and their small RNA profile suggest that their transcriptional activity is geared to produce piRNA precursors. Our results show a basal expression of NIRVS-derived primary piRNAs that are antisense to viral mRNA. These piRNAs could block novel infections with cognate viruses or they could interact with RNAi mechanisms to contain replication of incoming viruses at a level that does not become detrimental to mosquitoes. Albeit leading to opposite effects on vector competence, both mechanisms display functional similarites to the CRISPR-Cas system of prokaryotic adaptive immunity. Even if further studies are essential to clarify the effect of NIRVS-derived piRNAs on mosquito immunity, our study clearly demonstrates that *Ae. aegypti* and *Ae. albopictus* have a high number of NIRVS in their genome, which confers heritable immune signals.

The higher number of NIRVS in Aedeine than in Anophelinae mosquitoes correlates with competence for a larger number of arboviruses of Aedeine mosquitoes. In this regard, *Cu. quinquefasciatus* shows an interesting intermediate phenotype because it is phylogenetically closer to Aedeine mosquitoes, but vectors a smaller range of arboviruses than Aedeine mosquitoes and, like Anophelinae, it can vector more protozoans and nematodes than Aedeine [54]. Additionally, *Cu. quinquefasciatus* has a number of NIRVS and TE load comparable to Anopheline, but an expanded gene family like *Ae. aegypti* [2,55-56].

## CONCLUSIONS

NIRVS are regarded as viral fossils, occurring as occasional events during to the long co-evolutionary history of viruses and their hosts ([33,35]. The high abundance and diversity of NIRVS in the genomes of *Ae. aegypti* and *Ae. albopictus* and observation that NIRVS produce piRNAs and reside in piRNA clusters support the intriguing hypothesis that the formation and maintenance of NIRVS are coupled with the evolution of the PIWI pathway in these two species. This may have led to functional specialization of the expanded PIWI gene family, PIWI expression in the *soma*, and a role for the piRNA pathway in antiviral immunity [21,45]. This hypothesis is compatible with two scenarios. First, NIRVS formation is an occasional event, which occurs more frequently in Aedeine than Culicinae and Anopheline because of the higher abundance of retrotransposons in the genome of Aedeine mosquitoes [2]. NIRVS that have integrated by chance into piRNA clusters produce transcripts that are shuttled into the piRNA pathway. PIWI proteins loaded with viral sequences may target incoming viruses, possibly conferring selective advantage. Thus, an occasional event linked to a particular TE landscape may be the trigger for the functional specialization of PIWI proteins. This scenario remains compatible with the possibility that NIRVS outside of piRNA loci encode protein products that compete *in trans* with virus replication, thereby affecting vector competence [57]. Second, it has been hypothesized that PIWI proteins actively interact with incoming viruses and that they are loaded with episomal vDNAs and integrate them into piRNA clusters [58]. Under this scenario, the selective pressure favoring PIWI protein specialization would come primarily from viruses. Taken together our data show that the interaction between viruses and mosquitoes is a more dynamic process than previously thought and that this interplay can lead to heritable changes in mosquito genomes.

## METHODS

### *In silico* screening of viral integrations

Genome assemblies of *D. melanogaster* and 22 currently available mosquito species were screened *in silico* using tBLASTx and a library consisting of genome sequences of 424 non-retroviral RNA viruses and one DNA arbovirus (Additional File 1: Supplemental Table S1).

Tested mosquito species were classified in arboviral (*Aedes aegypti, Aedes albopictus, Culex quinquefasciatus*) and protozoan (*Anopheles gambiae, Anopheles albimanus, Anopheles arabiensis, Anopheles darling, Anopheles stephensi, Anopheles funestus, Anopheles atroparvus, Anopheles coluzzii, Anopheles culicifacies, Anopheles dirus, Anopheles epiroticus, Anopheles farauti, Anopheles maculatus, Anopheles melas, Anopheles merus, Anopheles minimus, Anopheles sinensis*) vectors depending on whether they most efficiently transmit arboviruses or protozoans to humans, respectively (Additional File 2: Supplemental Table S2). The non-vector *Anopheles christiy* and *Anopheles quadriannulatus* were also included in the analyses [59].

Host genome sequences of at least 100 bp and with high identity (e-values <0.0001) to viral queries were extracted from the respective insect genomes using custom scripts. When several queries mapped to the same genomic region, only the query with the highest score was retained. Blast hits were considered different when they mapped to genomic positions at least 100 bp apart from each other, otherwise they were included in the same NIRVS-locus.

All putative viral integrations were subjected to a three-step filtering process before being retained for further analyses to reduce the chance of false positives and ensure that the identified sequences are from non-retroviral RNA viruses [25]. Filtering steps included 1) a reverse-search against all nucleotide sequences in the NCBI database using the BLAST algorithm, 2) a search for ORFs encompassing viral proteins based on NCBI ORFfinder and 3) a functional annotation based on Argot^2^ [60].

While our search expanded the range of viral integrations identified in *Ae. albopictus* and *Ae. aegypti* [2,26,28], we cannot exclude that refinements of the current genome annotations of the species analyzed, especially in repeat regions, the application of alternative bioinformatic pipelines and the identification of novel viral species could lead to the characterization of additional integrations. Additionally, to reduce chance of false positives, our bioinformatics pipeline focused on sequences in which we could unambiguously identify viral ORFs, thus excluding viral sequences coming from UTRs or sub-genomic regions.

### Genomic data from 16 *Ae. albopictus* mosquitoes

Mosquitoes of the *Ae. albopictus* Foshan strain were used in this study. The strain was received from Dr Chen of the Southern Medical University of Guangzhou (China) in 2013. Since 2013, the Foshan strain has been reared in an insectary of the University of Pavia at 70-80% relative humidity, 28°C and with a 12-12 h light–dark photoperiod. Larvae are fed on a finely-ground fish food (Tetramin, Tetra Werke, Germany). A membrane feeding apparatus and commercially-available mutton blood is used for blood-feeding females.

DNA was extracted from single mosquitoes using the DNeasy Blood and Tissue Kit (Qiagen, Hilden Germany) following manufacturer’s protocol. DNA was shipped to the Polo D’Innovazione Genomica, Genetica e Biologia (Siena, Italy) for quality control, DNA-seq library preparation and sequencing on an Illumina HiSeq 2500. After quality control, retrieved sequences were aligned to the genome of *Ae. albopictus* reference Foshan strain (AaloF1 assembly) using the Burrows-Wheeler Aligner (BWA) [61] and marking identical read copies. The resulting indexed BAM files were used to calculate the counts of alignments, with mapping quality score above 10, which overlapped intervals of *Ae. albopictus* NIRVS using BEDTools [62]. Alignment files were visualized using the Integrative Genomics Viewer [63].

### Phylogenetic analyses

Deduced NIRVS protein sequences were aligned with subsets of corresponding proteins from *Flavivirus, Rhabdovirus, Reovirus* and *Bunyavirus* genomes using MUSCLE. Maximum likelihood (ML) phylogenies were estimated in MEGA6 [64], implementing in each case the best fitting substitution model. Statistical support for inferred tree nodes was assessed with 1000 bootstrap replicates. Figures were generated using FIGTREE (v.1.4) (http://tree.bio.ed.ac.uk/software/figtree/).

### Bioinformatic analyses of integration sites

Clustering of viral integrations in piRNA loci was estimated using cumulative binomial distribution, where the probability of integration was assumed to equal the fraction of the genome occupied by the respective genomic region. Genomic regions considered were piRNA clusters, coding regions and intergenic regions as previously defined [2,23,44]. A value of *P* < 0.05 suggests a statistically significant enrichment of these events in the corresponding genomic region (Table 2).

Analyses of TE enrichment in all non-retroviral integration sites as well as region 1 and region 2 of *Ae. aegypti* were based on RepeatMasker (version open-4.0.3, default parameters) using *A. aegypti* TEs retrieved from TEfam (http://tefam.biochem.vt.edu/tefam/), which was manually annotated. We used percent TE occupancy (percent of bases in the genomic sequence that match TEs) as an indication for possible enrichment of certain TEs. We did not use TE copy number as indications for TE enrichment because it is likely that some TEs can be broken into multiple fragments and be counted multiple times. We retrieved sequences of the viral integration sites plus 5 kb sequences flanking each side of the integration for the analysis. In addition, to identify potentially full-length TEs, 10 kb sequences flanking each side of the viral integration were analyzed by RepeatMasker (version open-4.0.3). Presence of full-length TEs was verified by comparing the length of masked sequences with the length of the annotated TEs.

### Analyses of piRNAs production from NIRVS

Small RNA deep-sequencing data of female *Ae. aegypti* (methoprene treated; SRX397102) [65] or *Ae. albopictus* mosquitoes (sugar-fed; SRX201600) [66] as well as PIWIs knockdown and IP libraries in Aag2 cells (SRA188616) [45] were downloaded from the European Nucleotide Archive. Subsequently, small RNA datasets were manipulated using the programs available in the Galaxy toolshed [67]. After removal of the 3’ adapter sequences, small RNAs were mapped to NIRVS sequences that were oriented in the direction of the predicted ORF, using Bowtie permitting one mismatch in a 32 nt seed [68]. From the mapped reads, size profiles were generated. For the analysis of nucleotide biases, the 25-30 nt reads were selected and separated based on the strand. The FASTA-converted sequences of small RNA reads were then trimmed to 25 nt and used as input for the Sequence-Logo generator (Galaxy version 0.4 based on Weblogo 3.3 [69]. piRNA counts on individual NIRVS were generated by mapping to NIRVS sequence after collapsing (near-) identical sequences (Additional File 5: Supplemental Table S5). Bowtie was used to map the small RNAs allowing one mismatch in a 32 nt seed. Only uniquely mapping reads were considered and the ‐‐best and the ‐‐ strata options were enabled. From the mapped reads, 25-30 nt small RNAs were selected. To identify secondary piRNAs, reads in sense orientation to viral ORFs that had an adenine at position 10 were selected. To avoid taking piRNAs into consideration that coincidentally contain a 10A, the population of 10A sense piRNAs was required to make up at least 50% of all sense piRNAs derived from the NIRVS. If this criterion was not met, sense reads from the corresponding NIRVS did not qualify as secondary piRNAs. Total piRNA counts and secondary piRNA counts were determined for F-NIRVS, R-NIRVS and *Reovirus* NIRVS and normalized to the corresponding library size. The size of the Ring-graph was scaled to reflect the total normalized read counts. piRNA counts on individual NIRVS was also determined from acetone treated female *Ae. aegypti*, male *Ae. aegypti*, blood-fed *Ae. albopictus* and male *Ae. albopictus* mosquitoes. The data was obtained from the same studies as described above.

To identify the PIWI dependency of NIRVS-derived piRNAs, we analyzed libraries from Aag2 cells transfected with double stranded RNA (dsRNA) targeting the somatic PIWI genes (Piwi4-6, Ago3) and a non-targeting control (dsRNA targeting luciferase, dsLuc) [45]. These datasets were mapped against the collapsed NIRVS dataset as described above. Since small RNA profiles were dominated by piRNA-sized reads, no further size selection was performed. The mean fold change in small RNA read counts was calculated for each PIWI knockdown condition compared to the negative control. To identify the PIWI proteins that NIRVS piRNAs associate with, we analyzed the IP libraries of PIWI proteins in Aag2 cells previously published in the same study. For the different PIWI IPs the enrichment of small RNA counts compared to a control GFP-IP was determined. Hierarchical clustering of NIRVS based on the combined fold changes of PIWI knockdowns and IPs was performed using multiple experiment viewer (Tm4). Clustering was based on Pearson correlation and performed independently for F-NIRVS and R-NIRVS.

### NIRVS transcriptional activity

RNA deep-sequencing data of *Ae. albopictus* and *Ae. aegypti* mosquitoes, including both DENV-infected and non-infected mosquitoes were downloaded from NCBI’s SRA. Libraries analyzed correspond to data SRA438038 for *Ae. albopictus, and* SRA058076, SRX253218, SRX253219 and SRX253220 for *Ae. aegypti*. RNA-seq reads were mapped using BWA [61] to NIRVS, after collapsing identical sequences (Additional File 5: Supplemental Table S5), and read counts were converted into RPKM using custom scripts.

To analyze AlbFlavi34 expression in different *Ae. albopictus* developmental stages, total RNA was extracted using Trizol (Life Technologies) from 3 pools of 5 entities for each condition (eggs, larvae, adult males, blood-fed and sugar-fed females). From each pool, a total of 100 ng of RNA was used for reverse transcription using the qScript cDNA SuperMix following manufacturer’s protocol (Quanta Biosciences). AlbFlavi34 expression was quantified in a 20 µL final reaction volume containing 10 µL of QuantiNova SYBR Green PCR Master Mix (Qiagen), 700 nM each forward (5’-CTTGCGACCCATGGTCTTCT-3’) and reverse (5’-GTCCTCGGCGCTGAATCATA-3’) primers and 5.0 µL cDNA sample on an Eppendorf RealPlex Real-Time PCR Detection System (Eppendorf). We used a two-step amplification protocol consisting of 40 cycles of amplification (95°C for 5 s, 60°C for 10 s) after an initial denaturation of 2 minutes at 95°C. AlbFlavi34 expression values were normalized to mRNA abundance levels of the *Ae. albopictus* Ribosomal Protein L34 (RPL34) gene [70]. QBASE+ software was used to visualise data and compare expression profiles across samples. Absence of *Flavivirus* infection was verified using a published protocol [24] on all samples before qRT-PCR.

## LIST OF ABBREVIATIONS

DENV: dengue viruses
ZKV: Zika virus
CHIKV: chikungunya virus
VEE: Venezuelan equine encephalitis
EEE: eastern equine encephalitis
LACV: La Crosse virus
RVFV: Rift Valley fever virus
WNV: West Nile virus
siRNA: small interfering RNA
piRNA: PIWI-interacting RNA
EVEs: Endogenous Viral Elements
NIRVS: Non-Retroviral Integrated RNA Viruses Sequences
N: Nucleoprotein
G: Glycoprotein
L: RNA-dependent RNA polymerase
ORF: Open Reading Frame
IP: immunoprecipitations
LTR: long terminal repeat (LTR) retrotransposons
TE: transposable elements
ISV: Insect Specific Virus
PTS: piRNA trigger sequence
BWA: Burrows-Wheeler Aligner
ML: Maximum likelihood
Ct=: cycle threshold
RPKM: Reads per kilobase per million mapped reads

## DECLARATIONS

### Ethics approval and consent to participate

Not applicable.

### Consent for publication

Not applicable

### Availability of data and material

The datasets used and/or analysed during the current study are available from the corresponding author on reasonable request.

### Competing interests

Not applicable.

### Funding

The project was supported by a European Research Council Consolidator Grant (ERC-CoG) under the European Union’s Seventh Framework Programme (grant number ERC CoG 615680) to R.P.v.R and an ERC-CoG under the European Union’s Horizon 2020 Programme (Grant Number ERC CoG 682394) to M.B. The funders had no role in study design, data collection and interpretation, or the decision to submit the work for publication.

### Authors’ Contributions

UP performed bioinformatic analyses and analysed the data; PM performed bioinformatic analyses, analysed the data and wrote the manuscript; RCL performed qRT-PCR analyses and analysed the data;, LO performed bioinformatic analyses and analysed the data; ER performed bioinformatic analyses and analysed the data; ZT performed bioinformatic analyses, analysed the data and wrote the manuscript; RvR, conceived the study and wrote the manuscript; MB conceived the study, analysed the data and wrote the manuscript.

## Acknowledgments

We thank Professor Anthony A. James (University of California at Irvine) for helpful comments on the manuscript. We thank Patrizia Chiari for insectary work.

